# In situ architecture and membrane fusion of SARS-CoV-2 Delta variant

**DOI:** 10.1101/2022.05.13.491759

**Authors:** Yutong Song, Hangping Yao, Nanping Wu, Jialu Xu, Zheyuan Zhang, Cheng Peng, Shibo Li, Weizheng Kong, Yong Chen, Miaojin Zhu, Jiaqi Wang, Danrong Shi, Chongchong Zhao, Xiangyun Lu, Martín Echavarría Galindo, Sai Li

## Abstract

Among the current five Variants of Concern, infections caused by the SARS-CoV-2 B.1.617.2 (Delta) variant are often associated with the greatest severity. Despite recent advances on the molecular basis of elevated pathogenicity using recombinant proteins, architecture of intact Delta virions remains veiled. Moreover, molecular evidences for the detailed mechanism of S-mediated membrane fusion are missing. Here we reported the in situ structure and distribution of S on the authentic Delta variant, and discovered invagination in the distinctive Delta architecture. We also captured fusion snapshots from the virus-virus fusion events, provided structural evidences for Delta’s attenuated dependency on cellular factors for fusion activation, and proposed a model of S-mediated membrane fusion. Site-specific glycan analysis revealed increased oligomannose-type glycosylation of native Delta S over that of the Wuhan-Hu-1 S. Together, these results disclose distinctive factors of Delta being the most virulent SARS-CoV-2 variant.

**In Brief:** Cryo-ET of intact SARS-CoV-2 Delta variant revealed its distinctive architecture and captured snapshots of its membrane fusion in action.

---

The continuing emergence of severe acute respiratory syndrome coronavirus 2 (SARS-CoV-2) variants imposes significant challenges for COVID-19 prevention and control. Among the five Variants of Concern (VOCs) designated by the World Health Organization (WHO), the B.1.617.2 (Delta) variant exhibits the highest pathogenicity. Hospitalized patients infected with the Delta variant showed severer symptoms and longer periods of illness compared to infections by other VOCs^1-3^. Epidemiologically, the Delta variant has 97% increased transmissibility^4^ and causes ∼1000 times of viral loads in patients^5^ than the ancestral Wuhan-Hu-1 (WT) strain.

The structural and functional impacts of spike (S) and nucleocapsid (N) mutations have been extensively studied. S is responsible for receptor binding and membrane fusion. Recent studies of the recombinant Delta S suggest that a group of mutations, including T19R, G142D, E156G, Δ157-158, L452R, T478K, D614G, P681R, and D950N, largely contributes to the elevated pathogenicity and transmissibility. Among these mutations, the T19R, G142D, E156G mutations and Δ157-158 locate at the N-terminal domain (NTD) antigenic supersite^6^. They cause dramatic structural rearrangement, and significantly impair the binding affinity of a large portion of NTD neutralizing antibodies (NAbs) to Delta S^6^. L452R and T478K locate at the receptor-binding domain (RBD) of S. Both mutations reinforce the S binding with the ACE2 receptor and L452R additionally increases the fusion efficacy by 50%^7, 8^. D614G is adopted by all five VOCs. It increases the proportion of RBD up conformations among prefusion S, leading to enhanced infectivity, and improves the fitness of S by reducing S1 shedding^9^. P681R locates at the furin cleavage site between S1/S2 and is unique to the B.1.617 lineage. It facilitates furin-mediated S1/S2 cleavage, significantly elevates and accelerates viral fusion, and enhances viral pathogenicity in hamsters^10^. Frequent mutations are also seen on N of all VOCs, which is responsible for packaging RNA into ribonucleoproteins (RNPs). Recruitment of RNPs by the membrane (M) protein triggers all viral components budding into the ER-Golgi intermediate compartment (ERGIC) to produce progeny virions^11^. In WT virions, loosely ordered RNPs attach to the interior of the viral envelope^12^ through M-N interactions^13^. Delta N-mutations include D63G, R203M, D377Y and in some clades additionally G215C and R385K. R203M improves viral assembly and increases the viral load by over 50-fold^14^, while G215C enhances N assembly with RNA^15^. Nevertheless, these impacts were concluded upon recombinantly expressed viral proteins. It remains unclear what the collective effects of these mutations are on the in situ structure and distribution of structural proteins, as well as the overall assembly of the authentic Delta virus.

The significantly enhanced fusogenicity is a hallmark of SARS-CoV-2 Delta S. It induced the most cell-cell fusion and has the highest fusion dynamics among S of the WT, D614G, Alpha, Beta^16^ and Omicron^17^ variants. Furthermore, multinucleated pneumocytes syncytium were present in the lungs of patients with severe COVID-19 symptoms^18, 19^. Such syncytia formation may support SARS-CoV-2 replication and transmission, immune evasion, and tissue damage. These in vitro/vivo evidences suggest that coronaviral fusogenicity may associate with its pathogenicity^16^. Despite the importance, in situ structural evidences on the detailed process of fusion have not been reported for coronaviral S, one of the largest known class-I fusion protein. For example, the specific conditions for fusion activation, the detailed sequence of coronaviral fusion, including structures of protein and membrane intermediates, the time point of RNP- and M-lattice disassociation, and the process of membrane remodeling have not been determined. Previous cryo-electron tomography (cryo-ET) studies on the influenza virus^20-22^, or Rift valley fever virus (RVFV)^23^-liposome mixture have captured intermediate steps of the glycoprotein-mediated fusion. Such observation on the Delta variant will be crucial not only in understanding the variant’s enhanced fusogenicity, but also providing a molecular basis for illustrating the SARS-CoV-2 fusion mechanism.

Here we combined cryo-ET, cryo-electron microscopy (cryo-EM) and mass-spectrometry (MS) for a comprehensive structural characterizations of intact Delta virions. We uncovered a dramatically distinctive architecture of Delta variant. We also determined the high-resolution structure, distribution, and glycan composition of Delta S in situ, which are compared with those of the previously characterized native WT^12^ or recombinant Delta S^9^. In addition, we captured snapshots of Delta virus-virus fusion “in action”, from which the sequence and intermediate steps of S-mediated membrane fusion were interpreted. These results provided molecular insights in membrane remodeling leading to SARS-CoV-2 fusion.

## Results

### Distinctive features of the Delta variant architecture

The SARS-CoV-2 Delta variant (hCoV-19/Hangzhou/ZJU-12/2021) was collected on May, 2021 from the sputum samples of a COVID-19 patient in Zhoushan, Zhejiang province, and was the first isolated Delta variant in China (GISAID: EPI_ISL_3127444.2). The clade lacks the E156G and Δ157-158 mutations on S. Virions were propagated in either Vero or Calu-3 cells in a BSL-3 laboratory and were fixed with paraformaldehyde (PFA) prior to concentration and cryo-electron microscopy (cryo-EM) analysis. The fixation has minor effects on protein structures at high-resolution and the overall assembly^12, 24, 25^. To avoid crushing the virions against the tube bottom during ultracentrifugation, we pelleted the virions onto the interface between 30% and 50% sucrose cushion.

With cryo-ET, we have imaged and reconstructed 1,032 Delta virions as volumetric data (Supplementary Table 1). The most distinctive feature of the Delta viral architecture is the presence of a large envelope concavity in the majority of virions (Fig. 1a-c, Supplementary Fig. 2f and Supplementary Movie 1). The concavity, which is rarely reported on enveloped viruses, is designated here as “invagination”. To validate that the invagination did not rise from cell lines that were used for virus propagation, or ultracentrifugation, we have imaged unconcentrated Delta virions from the infected Vero-cell supernatant (Supplementary Fig. 1a), concentrated Vero cell-propagated Delta virions (Fig. 1a) and concentrated Calu-3 cell-propagated Delta virions (Supplementary Fig. 1b). As a result, invagination is common in all conditions, but absent in Vero cell-propagated WT virions that were concentrated same as the above conditions (Supplementary Fig. 1c). For the convenience of description, key parts of the invaginated virions are named as in (Fig. 1f).

**Fig. 1.**
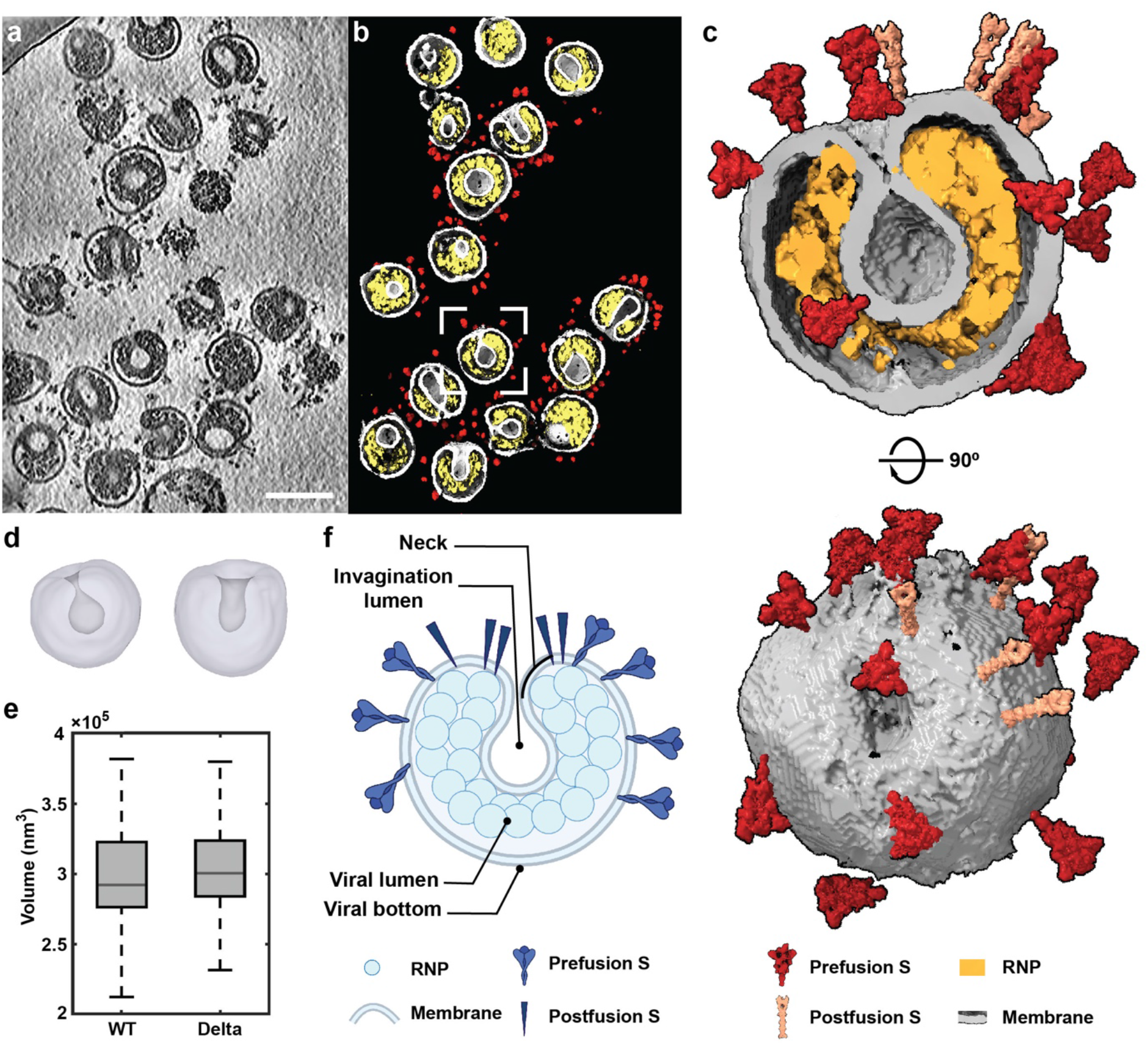
Molecular architecture of SARS-CoV-2 Delta variant. (**a**) representative tomogram slice (5 nm thick) showing pleomorphic SARS-CoV-2 Delta virions. Scale bar: 100 nm. (**b**) The tomogram in (a) was segmented to distinguish viral spikes (red), membrane (grey) and RNPs (yellow). Each virion in this tomogram harbors an invagination. (**c**) A topview and slice through a representative invaginated Delta virion showing the prefusion S (red), postfusion S (salmon), lipid envelope (grey) and RNPs (yellow). (**d**) Two representative types of invagination. One has a balloon-shaped lumen (left), and the other has a cylindrical shaped lumen (right). (**e**) Volume of WT and Delta virions. For invaginated virions, the volume does not include the invagination lumen. (**f**) Schematic of an invaginated virion with key parts of the virion named. Created with BioRender.

Scrutiny of the invaginated virions has revealed common features. First, invaginated virions are characterized by either a balloon-shaped, or a cylindrical lumen (Fig. 1d). These invaginations have an average depth of 45.8 nm, and connect the virion exterior through a neck with an average diameter of 7.6 nm (Supplementary Fig. 2e). Despite the invagination, Delta virions have similar volume within the viral lumen compared to WT virions (Fig. 1e). Second, S distribute only on the virion exterior, but rarely inside the invagination lumen. Third, the dramatic luminal reorganization is accompanied by the change of viral RNP-assembly. In WT virions, bead-like RNPs are often individually discernable, relatively homogeneously distributed, and hexagonally assembled on the cytoplasmic surface of the envelope^12^. While in invaginated Delta virions, the majority of RNPs form a dense layer wrapping around the invaginated envelope and leave the rest of viral lumen empty (Fig. 1a-c). Last, the viral envelope neighboring the invagination bottom (denoted as viral bottom) is largely devoid of RNPs and S, or sometimes even broken. Of the 721 invaginated virions, 216 have broken viral bottom. Lipid-bilayer densities from envelopes of the WT virions, Delta viral bottom and Delta invagination lumen were subtomogram averaged for comparison, revealing relatively thinner lipid bilayers on the Delta viral bottom (Supplementary Fig. 2g). This suggests that the area may have been subjected to extensive tension^26^.

### The molecular landscape of S on intact Delta virions

We combined cryo-ET and subtomogram averaging (STA) to investigate the assembly of Delta virions. In total we identified 22,695 prefusion S and 2,810 postfusion S from the tomograms, averaging 32 ± 12 S (24 ± 10 being prefusion S and 7 ± 4 being postfusion S) per virion. Remarkably, each Delta virion harbors on average ∼600% more postfusion S over WT virion, while the difference hardly exists for the prefusion S (Fig. 2e). Furthermore, STA and classification revealed that 36.4% prefusion S adopted the closed conformation, and 63.6% adopted the one RBD up conformation (Fig. 2d). S in two prefusion conformations, and the postfusion conformation, were reconstructed to 9.8, 13.0 and 11.9-Å resolution (Fig. 2a,b and Supplementary Fig. 3). Statistics of refined orientations of the prefusion S suggested freedom of S rotating around their stalks outside the envelope, with a mean tilting angle of 49° (std: 21.5°) relative to the normal axis of the envelope (Fig. 2b). For invaginated virions, S tend to accumulate around the neck area (Fig. 2f and Supplementary Fig. 2a-d). On envelope areas where RNPs are not attached to the cytoplasmic surface, there are neither prefusion S on the opposite surface (Fig. 1a-c, Supplementary Fig. 2a-d and Supplementary Movie 1). This cohabitation of prefusion S and RNPs suggests that the cytoplasmic region of S interact with N, or at least indirectly through M.

**Fig. 2.**
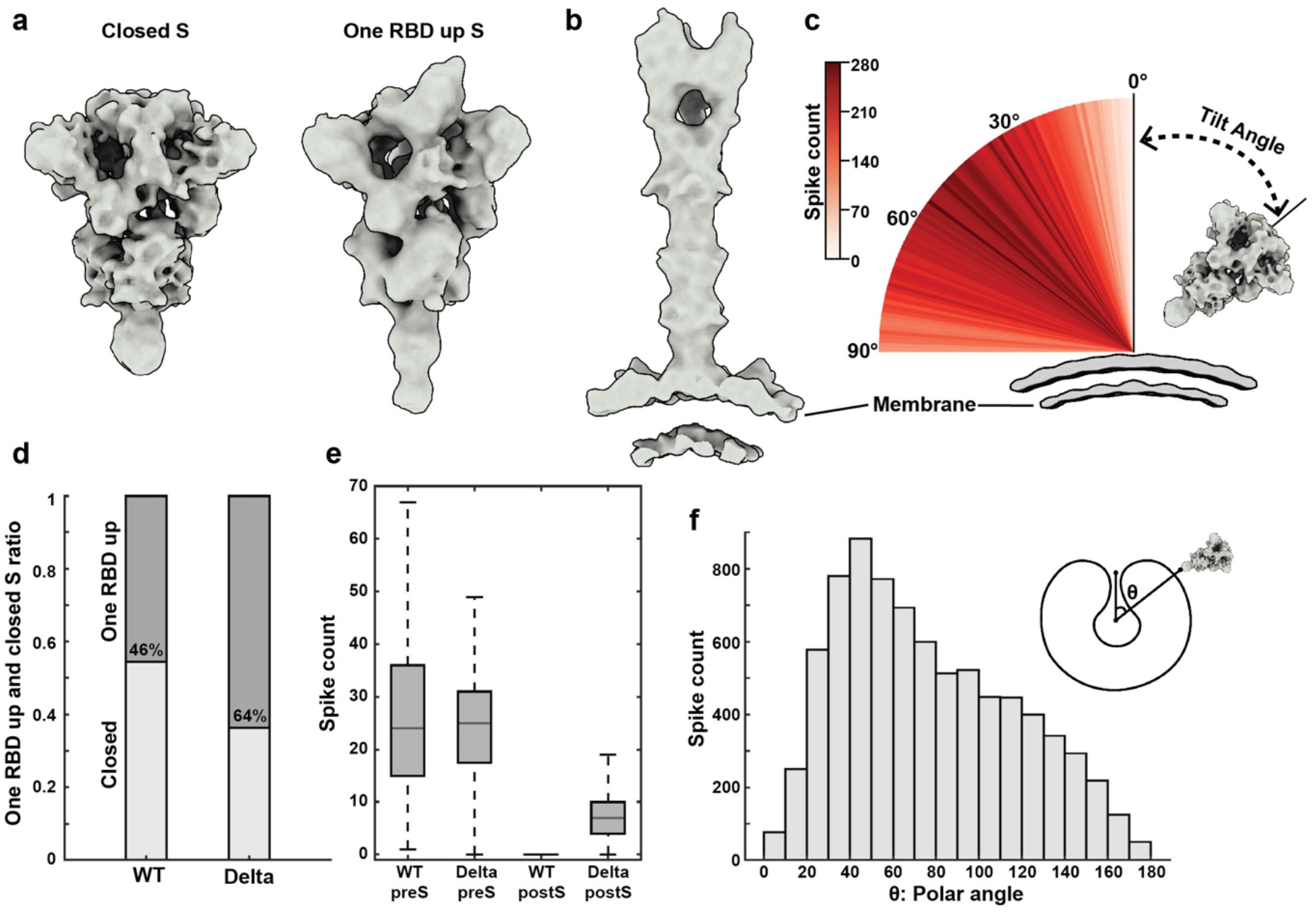
Native structures and distribution of S on the SARS-CoV-2 Delta virion surface. (**a**) The native structures of S in the closed and one RBD up prefusion conformations. (**b**) The native structure of S in the postfusion conformation. (**c**) Distribution of the spike tilt angle reveals a mean tilting angle of 49° (std: 21.5°) relative to the normal axis of the envelope. A representative S in the closed conformation on the envelope is shown. Step size: 1 degree. (**d**) Statistics on the percentage of one RBD up and closed S on WT and Delta virions. (**e**) Statistics on the average number of pre- and postfusion S per WT and Delta virions. (**f**) Statistics of S distribution on invaginated virions suggests that the spikes tend to accumulate around the neck area. As shown in the histogram, the invagination direction is defined by the vector from the virion center to the invagination neck; θ, or the spike latitude, defines the angle between the vector from the virion center to the spike stem and the invagination direction.

### The structure of S in situ on Delta virions and its glycan compositions

To gain further insights on the S structure in situ, S-trimers were imaged and refined by a single particle analysis (SPA) approach directly on the Delta viral surface (Supplementary Fig. 4 and Supplementary Table 2). Spikes were identified from the periphery of intact Delta virions, and aligned to a 4.39-Å resolution map of prefusion S. Focused classification on asymmetric RBD units revealed two classes, differing only in the density levels. Next, we separated and refined the three RBD closed S (55.3% of the data) and one RBD weak S (37.2% of the data) to 4.6- and 6.9-Å resolution, respectively (Supplementary Figs. 4 and 5). From the RBD closed S map, we built and refined an atomic model of S in situ on the viral surface (Fig. 3a). The model is compared to structures of recombinantly expressed WT S (PDB:6XR8)^27^ and Delta S (PDB:7SBK)^9^. Compared to the recombinant WT S, NTD of our structure has a clockwise rotation by 9.5°, when S2 of both structures are aligned (Supplementary Fig. 6a). Compared to the recombinant Delta S, three additional NTD loops (residues 1-26, 144-155, and 173-185), a receptor binding motif (RBM) loop (residues 468-489) and a CTD loop (residues 622-639, or the 630 loop) are disordered on our structure, reflecting the flexible nature of these sites in situ. The NTD in our structure extends 4.4° outward when S2 of both structures are aligned (Supplementary Fig. 6b). Moreover, residues 156-163 adopt a β-sheet conformation on our NTD while an α-helical conformation on that of 7SBK (Fig. 3a). This difference originates from the missing E156G and Δ157-158 mutations in our sample (Fig. 3d). The S2 structures are largely same between the in situ and recombinant Delta S: the S2’ cleavage sites are visible and uncleaved, and the furin cleavage sites are disordered. The only difference is on the stalk, with the C-terminus (terminates at E1151) of our structure nine residues shorter than 7SBK, reflecting the capability of spikes tilting around the highly flexible stalk on the virus (Figs. 2c and 3a).

**Fig. 3.**
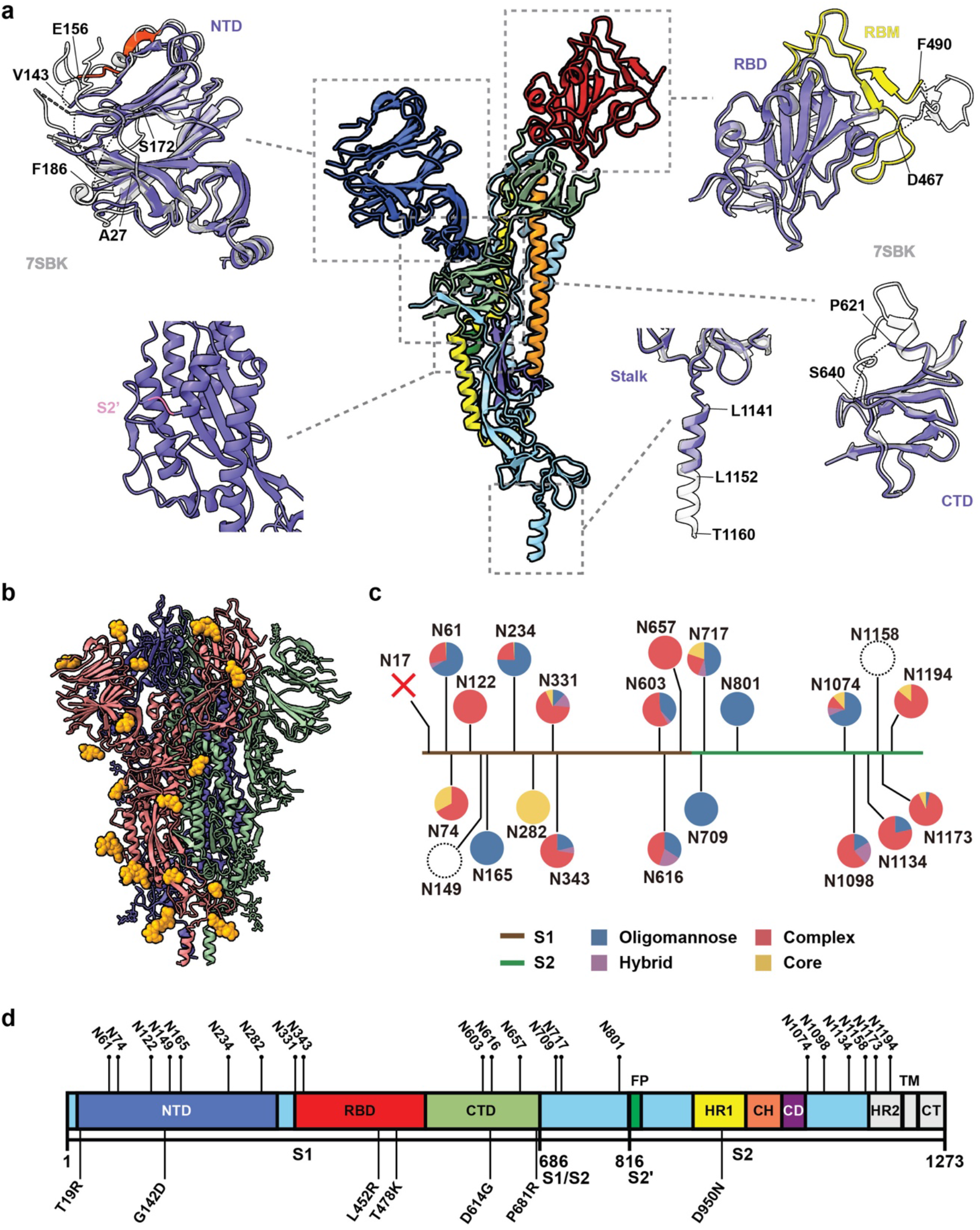
In situ structure and N-linked glycan compositions of SARS-CoV-2 Delta S. (**a**) In situ structural analysis of Delta prefusion S in closed conformation. The structure was determined to 4.39 Å resolution directly from the virion surface by single particle analysis (SPA). A protomer is colored by domains as shown in (d). Boxed regions including the RBD, NTD, CTD and stalk, are compared to those of the recombinant Delta S (PDB:7SBK, colored in gray). On the in situ structure, residues 156-163 adopt a β-sheet conformation on NTD (highlighted in red), three NTD loops (residues 1-26,144-155,173-185) are disordered; residues 468-489 are disordered on RBM, the rest of RBM is colored in yellow; residues 622-639 are disordered on CTD; the cleavage site on S2’ is in uncleaved state; the stalk is ordered until residue L1152. (**b**) Glycosylation profile of the in situ Delta S structure, N-linked glycans are colored in orange. (**c**) The identity and proportion of the 19 N-linked glycans on the native full-length Delta S were analyzed by MS and are shown in pie charts. The missing glycan N17 due to the mutation was shown as a cross. The undetected glycans N149 and N1158 were shown as dashed circles. (**d**) Sequence schematic of Delta S, showing the mutation sites and glycosylation sites. NTD, N-terminal domain; RBD, receptor binding domain; CTD, C-terminal domain; S1/S2, S1/S2 cleavage site; S2’, S2’ cleavage site; FP, fusion peptide; HR1, heptad repeat 1; CH, central helix; CD, connector domain; HR2, heptad repeat 2; TM, transmembrane anchor; CT, cytoplasmic tail.

The T19R mutation is known to abrogate the N17 glycan. Among the remaining 21 N-linked glycosylation sites, we are able to build 16 glycans from their densities (Fig. 3b). The remaining five glycans locate on disordered NTD loops or the unsolved stalk region. We further performed a site-specific glycan analysis of the native Delta S by resolving the virus sample on SDS-PAGE, and analyzing the bands corresponding to S1 and S2 with mass spectrometry (MS). Of the 19 glycans detected and analyzed, the native Delta S glycans contain more oligomannose-type glycosylation compared to the native WT S glycans^12, 28^ (Fig. 3c). N149 and N1158 are missing in both the cryo-EM structure and MS results.

### Visualization of Delta S-mediated membrane fusion

S-involved membrane remodeling, including membrane insertion, dimpling or pinching of membranes, were frequently observed in tomograms of purified Delta virions. These observations are reminiscent of typical intermediate steps of class-I fusogen mediated membrane fusion^29^. We sequenced these observations into six steps of a fusion process (Fig. 4a, b and Supplementary Fig. 7), and further characterized these steps to illustrate the SARS-CoV-2 fusion mechanism (Fig. 4c, d): 1) Two envelopes connected by a long, thin and straight bar density, which has an average length of 26.3 nm, 34% longer than the length of a postfusion S. This is denoted S2-mediated bridging. 2) Two envelopes connected by long, thin but kinked bars, which consist of a shorter rear arm (average length: 8.9 nm) and longer forearm (average length: 15.4 nm) linked by an elbow region, with an average kink angle of 143°. This is denoted S-mediated dimpling. 3) Two “kissing” envelopes featuring membranes in primary contact. Thin, straight bars were often seen surrounding the contact area. 15 bars from 8 independent events were averaged by STA, which reconstructed a 21.7 nm long spike inserted between two envelopes. Fitting the spike with a postfusion S structure (PDB:6XRA) suggests that it is close to the postfusion form. Segmentation of one event revealed a group of six spikes radiating away from the membrane contact point, forming a star-shaped rosette. This is similar to influenza HA mediated pinching^22^. During the three steps, RNPs remain attached to the cytoplasmic side of the membrane underlying the participating spikes, suggesting that the N-associated M-lattice has not become disrupted yet^13^. 4) Two envelopes forming a tightly docked interface. The interface is formed by two bilayers, as measured by the density profile, and can extend as wide as 36 nm. At least one involved virion is distorted, with its membrane pulled away from its RNPs towards the interface, suggesting the M-lattice has become disrupted. 5) Two envelopes with their membranes forming a single-bilayer interface, as measured by the density profile. This is denoted hemifusion. 6) One large virion harboring S on the envelope while containing two sets of RNPs inside. This is denoted fusion completion. The above phenomena were very rare in our previous data of WT SARS-CoV-2^12^.

**Fig. 4.**
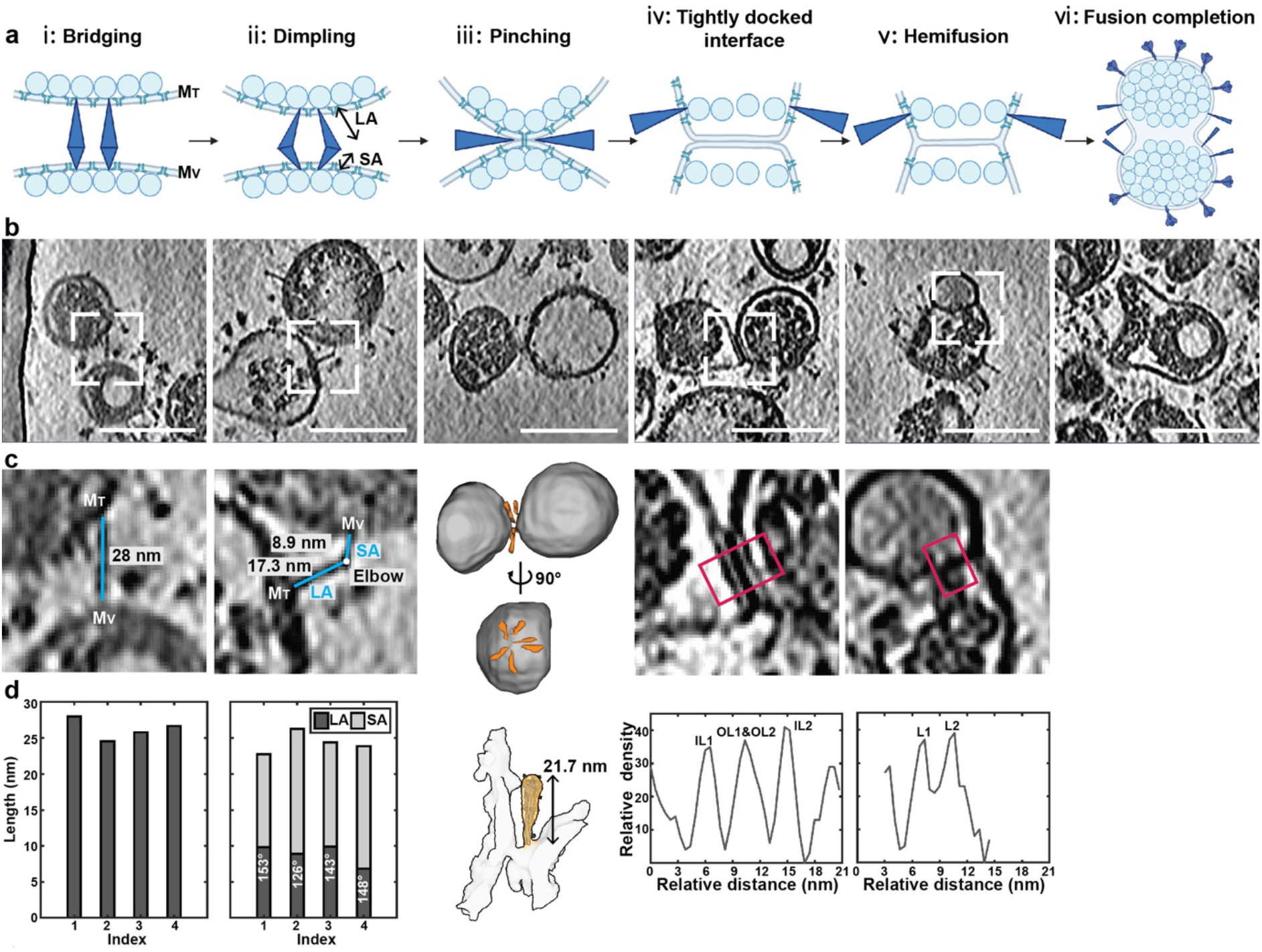
Sequence of membrane fusion of SARS-CoV-2 Delta variant. (**a**) Observations of intermediate membrane fusion steps were generalized into six stages: i: bridging; ii: dimpling; iii: pinching; iv: tightly docked interface; v: hemifusion and vi: fusion completion. Created with BioRender. (**b**) Representative tomogram slices (thickness 5 nm, scale bar 100 nm) of the corresponding six fusion stages. (**c-d**) Characterization of the fusion steps. For the bridging stage, lengths of the bridging spikes were measured for four individual events. For the dimpling stage, measurements of the angle between and length of the shorter arm (SA) and longer arm (LA) of the mediating spikes for four individual events were shown. For the pinching stage, the tomogram in (b) was segmented. The relative position of the spikes (orange) and the envelope (grey) was shown from the side- and topview. Subtomogram averaging of 15 spikes from 8 independent pinching events shows an S (orange) pinching the envelopes (grey) of two virions. A postfusion S (PDB: 6XRA) was fitted to the map for comparison. For the tightly docked interface and hemifusion stages, density profiles across the interface (red box) were plot, showing two lipid bilayers (IL: inner leaflet, OL: outer leaflet) for the tightly docked interface and one lipid bilayer (L: leaflet) for the hemifusion stage.

## Discussion

In this work, the in situ analysis of S provided molecular basis for the enhanced infectivity and immune evasion of the Delta variant verses the WT strain on multiple scales. First, the Delta variant harbors 23.1% more S than the WT strain. Given their similar average diameters, the population density of S is higher on the Delta variant. Second, the Delta prefusion S rotate more vigorously around their stalks on the envelope. Third, 18% more prefusion S adopt the one RBD up conformation. Last, the disordered NTD loops (residues 1-26, 144-155, 173-185 and 245-262) on our structure reside largely on the vulnerable NTD antigenic supersite, reflecting the selective pressure on Delta variant for immune evasion. In terms of glycan composition, there is a greater proportion of oligomannose on Delta S than that on WT S. This implicates that the significantly higher viral production of Delta variant in the cell outspeeds the capacity of the cell to process the large number of glycan sites present on S in its Golgi. The increased oligomannose on Delta S may also facilitate virus-cell attachment and viral entry through oligomannose-GlcNAc glycan-glycan interactions, as was observed on HIV^30^. The structure, glycan analysis and statistics of S on virions also provide in situ profiles of effective antigens, which is an important reference for vaccine development.

The virus-virus fusion observed in Delta virions could rises from various factors. First, Delta virions are more pleomorphically assembled compared to WT virions. Apart from being invaginated, Delta virions have large areas of bald envelope and much less homogenously distributed RNPs to support the viral assembly. Another possibility is its attenuated dependency on cellular factors for fusion activation. Angiotensin converting enzyme 2 (ACE2)-binding followed by cleavage of S2’ site either by transmembrane serine protease 2 (TMPRSS2), or by cathepsin L have been suggested as necessary steps for priming the fusion activation^31^. However, postfusion S have been found on intact SARS-CoV-2 WT^12, 25, 32^ and D614G^24^ virions, implying the existence of an uncharacterized, cellular-factor independent route of fusion activation that leads to S1-disassociation and S2-refolding. This alternative route of fusion activation may be enhanced by characteristic mutations on Delta S, such as L452R and P681R^7, 10^, as recent studies showed syncytia can be induced either by Delta S on cells with no exogenous ACE2^9^ or by D614G/P681R S on cells with no exogenous TMPRSS2^10^. Our observations of the significantly more postfusion S per Delta virion, and the virus-virus fusion provided direct molecular evidences for the spontaneous fusion activation. Besides the L452R and P681R mutations, this attenuated dependency also originates from increased instability of S caused by the disordered 630 loop (residues 622-639) discovered on our in situ structure. Ordered 630 loops insert into the NTD and CTD, help stabilizing their relative position and prevent S1 shedding. However, in our structure the 630 loops are disordered, with NTD and CTD forming a larger wedge than 7SBK (Supplementary Fig. 6b). The observation suggests that dislocation of the 630 loop from CTD destabilizes this domain and allows S1 shedding. Nevertheless, despite the possibility of spontaneous fusion activation, given the dominant population of prefusion S on intact virions, and the uncleaved S2’ site observed on the in situ structure, the cellular-factor dependent fusion activation remain as the dominant routes for viral fusion and entry.

We also exploited the virus-virus fusion present in Delta variant as an opportunity to elucidate the SARS-CoV-2 fusion process. For the virus-liposome fusion, where the unsupported liposome membrane is clearly pulled towards the virus^20^, it is unambiguous to distinguish the viral membrane (M_V_) from the target membrane (M_T_). However, this is not obvious for the virus-virus fusion, since both sides of the membranes are supported by M- and RNP-lattices. Comparison of S2 in its prefusion (PDB:7SBK) and postfusion (PDB:6XRA) conformations suggests that the longer arm of the dimpling S2 comprises the fusion peptide (FP), the heptad repeat 1 (HR1) and the central helix (CH) domain; and the shorter arm comprises the heptad repeat 2 (HR2) and the transmembrane anchor (TM) domain. The two arms are linked by an elbow region comprised of a relatively rigid β-rich module (residues 1035-1140). These identifications are supported by the averaged length and angles measured from the dimpling S2 (Fig. 4d), and helped us determined the shorter-arm-attached membrane to be M_V_ and the other to be M_T_. As fusion proceeds, at least one involving virion becomes distorted, with its membrane pulled away from its RNPs. This membrane area is often devoid of spikes, suggesting that the pulling has possibly disrupted the M-lattice, whose interaction with S and N plays key roles in viral assembly^33^. Among the six steps of fusion activities present in the tomograms, the pinching and tightly docked interface were the most frequently observed events, suggesting that these steps are metastable. The hemifusion was relatively rare, which reflects its transient nature. These observations are in agreement with the Influenza A virus hemagglutinin (HA) mediate fusion^20-22^. Upon these observations, we proposed a model for the SARS-CoV-2 S-mediated spontaneous fusion (Fig. 5). To begin with, S1 spontaneously sheds. Then HR1 unfolds and inserts FP into the target membrane, forming an extended intermediate. The membrane-embedded M-lattice and the M-affiliated RNP-lattices remain intact at this stage. The FP-HR1-CH region then refolds around the β-module towards the HR2-TM region, pinching the viral and target membrane. Since this stage, the M-lattice has become disrupted and the RNP-lattice has disassembled from the viral membrane. After going through the tightly docked and hemifusion transitions, the pinched membranes finally fuse. The exhausted S2 eventually adopts a postfusion conformation. A necessary condition for the virus-virus fusion is the proximity of virions. Since our virions were fixed prior to concentration, we argue that the fusion most likely occurred during virus budding or egressing. In both steps, the assembling or assembled SARS-CoV-2 virions can accumulate in small cellular compartments, such as ER-Golgi intermediate compartment (ERGIC)^11, 34^ or lysosomes^11, 35^.

**Fig. 5.**
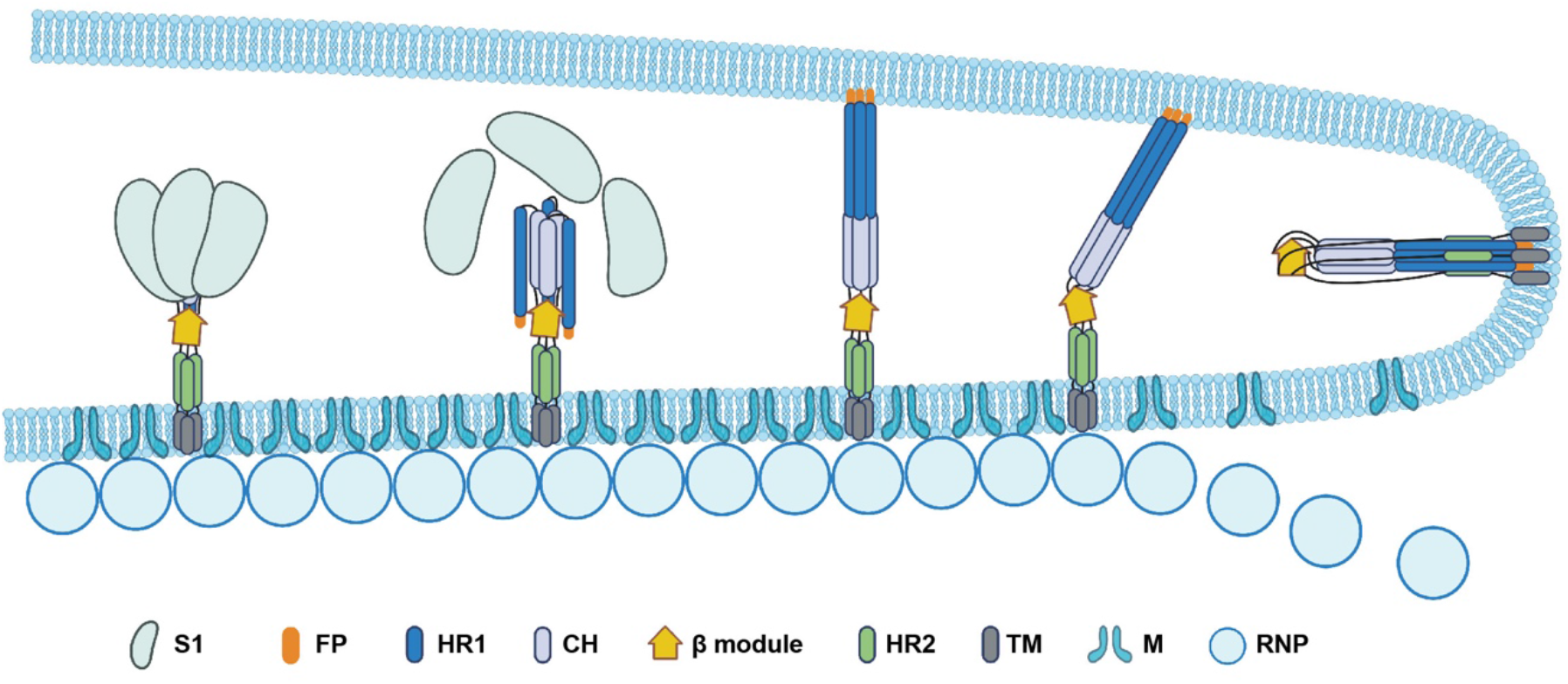
Proposed model for SARS-CoV-2 S-mediated membrane fusion. Model of SARS-CoV-2 S-mediated spontaneous membrane fusion. S1 spontaneously sheds, then HR1 unfolds and inserts FP into the target membrane, forming an extended intermediate. The FP-HR1-CH region then refolds around the β-module towards the HR2-TM region, pinching the viral- and target membrane. Since this stage, the M- and RNP-lattices have become disrupted and disassembled from the viral membrane. After fusion completion, S2 adopts a postfusion conformation. HR1, heptad repeat 1; CH, central helix; FP, fusion peptide; β module, a β-rich domain (residues 1035-1140); HR2, heptad repeat 2; TM, transmembrane anchor; M, membrane protein; RNP: ribonucleoprotein. Created with BioRender.

The unexpected invagination is another distinctive feature of Delta variant. The origin and mechanism of invagination formation remain unclear. We hypothesized that the invagination most likely originated from the N-mutations. Recent studies suggest that single N-mutations on R203 or G215 significantly elevate the infectivity, viral assembly efficiency, or virulence, possibly through enhanced N assembly with viral RNA and alteration of N-oligomerization on RNPs^14, 15, 36^. The structure and assembly of N is highly plastic due to the disordered linkers present near the N-terminal RNA binding domain (N-NTD) and the C-terminal dimerization domain (N-CTD). Among the VOCs, most hotspots of N-mutations locate on these disordered linkers, implying that SARS-CoV-2 might exploit these mutations to achieve greater efficiency of assembly. The reformation of RNP-structure and assembly observed on Delta virions (Fig. 1a-c and Supplementary Fig. 1) suggests possibility of an altered mode of viral budding into ERGIC (Supplementary Fig. 8). In this model, RNPs are recruited by the membrane-residing M of an EGRIC, and bud into the ERGIC while forming a dimple. Continuing budding enlarges the dimple into an invagination. With increased budding efficiency or change of RNP-assembly due to N-mutations, the viral bottom was left either bald or ruptured after budding completion. We have observed similar invagination on another beta-coronavirus (unpublished data), suggesting that the invagination may be a common feature on a broad range of beta-coronaviruses.

In conclusion, in situ structural evidences on multiple scales have collectively suggested that Delta S are tuned in a more unstable state, attenuating its dependency on cellular factors for fusion. This is alarming for strategies that tried to use TMPRSS2/cathepsins inhibitor to block virus entry^37^. Furthermore, comparison of the Delta and Omicron variants with the WT strain suggested that the fusogenicity of a SARS-CoV-2 strain may associate with its pathogenicity. Thus, the prominent fusogenicity may explain for the greater severity and unusual symptoms of Delta infections^10, 17^, and is alerting for future variants that may carry similar fusogenicity-enhancing mutations. Molecular evidences on the altered mode of budding into ERGIC by Delta variant, as well as why this significantly increases the viral assembly efficiency are of great interests for future studies.

## Methods

### Date and materials availability

Electron microscopy maps and the tomogram in Figure 1a have been deposited in the Electron Microscopy Data Bank under accession codes EMD-32205, EMD-32206, EMD-32207, EMD-32208 and EMD-33291. The atomic structure coordinates are deposited in the RCSB Protein Data Bank (PDB) under the accession XXXX. Viral sequence has been deposited in GISAID under accession ID: EPI_ISL_3127444.2. The mass spectrometry proteomics data have been deposited to the ProteomeXchange Consortium via the PRIDE^38^ partner repository with the dataset identifier PXD033342.

### Sample preparation

SARS-CoV-2 Delta variant was isolated from sputum samples of COVID-19 patient. The virions were propagated in Vero cells (ATCC CCL-81) or Calu-3 cells (ATCC HTB-55). Sputum was diluted by 5 volumes of Modified Eagle Medium (MEM) complete medium supplemented with 2% fetal bovine serum (FBS), Amphotericin B (100 ng/ml), Penicillin G (200 units/ml), Streptomycin (200 μg/ml) and centrifuged to remove impurities at 3000 rpm for 10 min in room temperature. Finally, the supernatant was collected and filtered through a 0.45 μm filter. 3 ml of filtered supernatant was added to Vero cells in a T25 culture flask. After incubation at 35 °C for 2 hours to allow binding, the inoculum was removed and replaced with 6ml fresh culture medium. The cells were incubated at 35 °C and observed daily to evaluate cytopathic effects (CPE). When CPE was present in most cells, the SARS-CoV-2 in culture supernatant was detected by qRT-PCR, sequencing and immunofluorescence. The viral genome sequence was uploaded to the GISAID with the ID of hCoV-19/Hangzhou/ZJU-012/2021 Vero EPI_ISL_3127444.2. For the preparation of enough virus samples, viruses were proliferated using Vero cells or Calu-3 cells in T75 culture flasks. On four days post-infection, 100 ml cell supernatant was cleared from cell debris at 4,000 g centrifugation for 30 min and inactivated with paraformaldehyde (PFA; final concentration 3%) for 48 hours at 4 °C. The inactivated virus samples were validated using cell infection and immunofluorescence staining. The supernatant was kept at 4 °C afterwards. All experiments involving infectious virus were conducted in approved biosafety level (BSL)-3 laboratory in the First Affiliated Hospital of Zhejiang University. SARS-CoV-2 Wuhan-Hu-1strain was processed as described in ^12^.

Purification, concentration, biochemical analysis and sample preparation for electron microscopy of inactivated virions were carried out in a BSL-2 lab at Tsinghua University. For cryo-ET, discontinuous sucrose cushion was applied. The virion was sedimented twice through a 30% sucrose band to 50% band in PBS buffer by ultracentrifugation (Beckman, IN) at 100,000 g, 4 °C for 3-5 hours. The solution at the sucrose interface were collected. For cryo-EM, the fixed virions were pelleted through 30% sucrose cushion by ultracentrifugation at 100,000 g, 4 °C for 3 hours. The pellet was resuspended in 60 µl PBS buffer.

### Mass spectrometric analysis

50 µg purified virion sample were heated to 100 °C for 15 min and characterized by 4 to 12% NuPAGE™ Bis-Tris gel (Invitrogen, Carlsbad, CA). The protein band was stained by One-Step Blue Protein Gel Stain (Biotium, Fremont, CA).

For glycan analysis, two replicates of gel bands corresponding to S protein were excised from the gel, reduced with 5 mM of DTT and alkylated with 11 mM iodoacetamide which was followed by in-gel digestion with trypsin (Promega, Madison, WI), chymotrypsin (Promega, Madison, WI) or alpha lytic protease (Sigma-Aldrich, St. Louis, MO) in 50 mM ammonium bicarbonate at 37 °C overnight. The sample was quenched by adding 10% trifluoroacetic acid (TFA) to adjust the pH to below 2. The peptides were extracted twice with 0.1% TFA in 50% acetonitrile aqueous solution for 1 hour and then dried in a speedVac. Peptides were dissolved in 25 μl 0.1% trifluoroacetic acid and 6 μl of the extracted peptides was analyzed by Orbitrap Q Exactive HF-X (Thermo Fisher Scientific, Bremen, Germany).

For LC-MS/MS analysis, the peptides were separated by a 120 min gradient elution at a flow rate 0.30 µl/min with a Thermo-Dionex Ultimate 3000 HPLC system, which was directly interfaced with an Orbitrap Q Exactive HF-X mass spectrometer (Thermo Fisher Scientific, Bremen, Germany). The analytical column was a home-made fused silica capillary column (75 µm ID, 150 mm length; Upchurch, Oak Harbor, WA) packed with C-18 resin (300 Å, 5 µm, Varian, Lexington, MA). Mobile phase A consisted of 0.1% formic acid, and mobile phase B consisted of 100% acetonitrile and 0.1% formic acid. An Orbitrap mass spectrometer was operated in the data-dependent acquisition mode using Xcalibur 4.3.73.11 software and there was a single full-scan mass spectrum in the Orbitrap (300–1800 m/z, 60000 resolution) followed by 3 s data-dependent MS/MS scans in an Ion Routing Multipole at stepped 27, 30, 33 normalized collision energy (HCD).

Glycopeptide fragmentation data were extracted from the raw file using Byonic™ (Version 2.8.2). The MS data was searched using the Protein Metrics 309 N-glycan library. The search criteria were as follows: Non-specificity; carbamidomethylation (C) was set as the fixed modifications; the oxidation (M) was set as the variable modification; precursor ion mass tolerances were set at 20 ppm for all MS acquired in an orbitrap mass analyzer; and the fragment ion mass tolerances were set at 0.02 Da for all MS2 spectra acquired.

Same as ^12^, the intensities of the same glycopepide in each site were combined and analyzed for proportion. Data with score under 30 and abnormal value with intensity above 1×10^10^ was removed. The glycans were classfied into oligomannose, hybrid, complex and core type based on composition. Hybrid and complex type glycan were subdivided according to fucose component and antenna. Mean of the two replicates represents ratio of each glycan type.

### Cryo-electron tomography and electron microscopy

For cryo-electron tomography, 10 µl sample was applied onto a glow discharged copper grid coated with holey carbon (R 2/2; Quantifoil, Jena, Germany), and subsequently dipped onto 500 µl deionized H_2_O for 1 second to clear the sucrose. A drop of 3 µl gold fiducial beads (10 nm diameter; Aurion, The Netherlands) was applied and the grid was blotted for 4.5 s, vitrified by plunge-freezing into liquid ethane using a Cryo-plunger 3 (Gatan, CA). The grids were imaged on a Titan Krios microscope (Thermo Fisher Scientific, Hillsboro, OR) operated at a voltage of 300 kV equipped with a GIF Quantum energy filter (slit width 20 eV) and K3 direct electron detector (Gatan, CA). Virions were recorded in super-resolution mode at a nominal magnification of 64,000×, resulting in a calibrated pixel size of 0.68 Å. 150 sets of tilt-series data were collected using the dose-symmetric scheme^39^ from -60° to 60° at 3° steps and at various defocus between -2.0 and -4.0 µm in SerialEM^40^. At each tilt, a movie consisting of 8 frames was recorded with 0.0265 s/frame exposure, giving a total dose of 131.2 e^-^/Å^2^ per tilt series.

For single particle analysis, the sample was prepared similarly to the cryo-ET protocol, except for using copper grid coated with holey carbon (R 1.2/1.3; Quantifoil, Jena, Germany), applying 4 µl virus sample and not adding fiducial beads. The grids were imaged on the same equipment as cryo-ET. Movies of micrographs were collected using AutoEMation 2 (written by Jianlin Lei) under super resolution mode at a nominal magnification of 81,000×. The pixel size was 0.541 Å/pixel. Each movie consists of 48 frames and was recorded using a total dose of 50 e^-^/Å^2^. In total, 25,851 movies were collected at various defocus between -1.0 and -3.0 µm.

### Cryo-electron tomography data processing

Tilt series data was analysed in a high-throughput pre-processing suite^12^ developed in our lab. The electron beam induced motion was corrected using MotionCor2^41^ by averaging eight frames for each tilt. Defocuses of the tilt series were measured using CTFFIND4^42^. The tilt series were contrast transfer function corrected using Novactf^43^, 150 tilt-series with good fiducial alignment and relative thin ice thickness were reconstructed to tomograms by weighted back projection in IMOD^44^, resulting in a final pixel size of 1.36 Å/pixel. The tomograms were 2 ×, 4 × and 8 × binned for subsequent processing. The 8 × binned tomograms were further processed by IsoNet^45^ to compensate for the missing wedge and enhance the contrast.

To characterize the viral morphologies in 3D, we segmented 276 intact invaginated virions from the IsoNet corrected 8× binned tomograms using 3Dslicer^46^, and measured dimensions of the invagination. To characterize the viral membrane fusion in 3D, we measured the angle and length of dimpling stage spikes from the IsoNet corrected tomograms in 3DSlicer. For the pinching stage spikes, 15 manually picked spikes were extracted into boxes of 64×64×64 voxels from IsoNet corrected tomograms and aligned by subtomogram averaging using Dynamo. The density profile of membranes at the tightly docked interface and hemifusion stages were measured in IMOD from the 30 Å low-passed, 4 × binned tomograms.

Particle identification was carried out using ilastik^47^. 23,223 prefusion S were automatically segmented and 2,810 postfusion S were manually segmented in ilastik. The coordinates of identified particles were calculated from the output files, and were transformed into Dynamo-readable format. The initial spike orientations were estimated from vectors normal to the local membrane.

Subtomogram averaging was done using Dynamo^48^ following a process we established earlier^12^. For the prefusion S reconstruction, 23,223 subtomograms were extracted from 8 × binned tomograms into boxes of 64×64×64 voxels and EMD-30426^12^ was used as template for their alignment. After removing 528 overlapping particles, the remaining 22,695 particles were extracted from 4 × binned tomograms into boxes of 96×96×96 voxels for further alignment. The resolution was restricted to 30 Å and C3 symmetry was applied at this stage. Subsequently, the particles were subjected to multi-reference alignment imposing C1 symmetry using EMD-30426 and EMD-30427 lowpassed to 30 Å resolution as the templates, resulting in 8,269 spikes (36.4%) classified into closed conformation and 14,426 spikes (63.6%) into one RBD up conformation. Coordinates of the two spike conformations were used to extract boxes of 160×160×160 voxels from the 2 × binned tomograms for further alignment. To prevent overfitting, a customized ‘gold-standard adaptive bandpass filter’ method^12^ was used for the alignment at this stage, and a criterion of 0.143 for the Fourier shell correlation were used to estimate the resolution. The 2 × binned spikes in the closed and one RBD up conformations were independently further aligned imposing C3 or C1 symmetry respectively, to 9.8 and 13.0 Å resolution. The prefusion S maps were lowpassed according to the estimated local resolutions of the reconstructed subunits. Universal empirical B-factors in the range of -1,500 ∼ -2,000 were applied to sharpen the closed S.

For the postfusion S reconstruction, 2,810 subtomograms were extracted from 4 × binned tomograms into boxes of 96×96×96 voxels, which were aligned using EMD-30428^12^ as the template. The resolution was restricted to 30 Å and C3 symmetry was applied at this stage. The refined coordinates were used to extract postfusion S from the 2 × binned tomograms into boxes of 160×160×160 voxels for gold-standard alignment. Subsequent alignment achieved 11.9 Å resolution.

### Cryo-EM data processing

Micrographs were motion-corrected and dose-weighted using RELION implementation of MotionCor2^41^. Subsequently, non-dose-weighted sums of power spectra were used to estimate the CTF with CTFFIND4^42^. Initial micrographs were deconvolved using Warp^49^, and 7,950 spikes were manually picked from 474 micrographs for training Topaz neural network. The trained Topaz neural network^50^ was used to automatically pick the spikes. 674,792 particles were auto-picked by Topaz and extracted in bin 4. Single particle analysis was done using RELION-3.1.0^51^. 3D classification was performed using EMD-21452 low-pass-filtered to 30 Å as the reference to remove the junk particles and revise the initial coordinates of auto-picked particles. 105,707 particles with revised coordinates were re-extracted in original scale. Two more rounds of 3D classification were performed using reference reconstructed from this dataset. The classification was ended when the RBD density were complete and no improvement for further 3D classification. The consensus class at this stage has 45,232 particles. This class was further subjected to Bayesian polishing and CTF refinement, yielding a RBD closed S-trimer at 4.39 Å resolution. The map was locally filtered and used for model building. This class was symmetry expanded and 135,756 asymmetric RBD densities were subtracted for local classification to sort different conformations. 23,842 RBD units showed missing RBM loop compared to the other 111,894 RBD units showed complete RBD density. 25,001 RBD closed S, 16,846 one RBD weak S, 3219 two RBDs weak S, and 186 three RBD weak S were found after the local classification. The RBD closed S were further refined to 4.6 Å resolution with C3 symmetry imposed. The one RBD weak S were further refined to 6.9 Å resolution with C1 symmetry imposed. The last two classes did not have enough particles for 3D reconstruction.

### Model building and refinement

The initial models used for model building are WT SARS-CoV-2 NTD (PDB:6XR8 from residue 13 to 304) and Delta SARS-CoV-2 trimer (PDB:7SBK from residue 305 to 1160) due to the lack of mutations and deletions on residues 156-158 on our sample. The two separate models were merged and manually adjusted in Coot. Loops 70-76, loop 144-155, loop 173-185, loop 245-262, loop 468-489, loop 622-639 that have not been solved in our map were manually deleted in Coot. Glycans were manually added in 16 solved glycosylation sites. G142D were manually added. Then the model was real-space refined in PHENIX. Steric clashes and sidechain rotamer conformations were improved manually in Coot after refinement. The final model is validated in PHENIX. The geometry and statistics are recorded in Extended Data Supplementary Table 2. The unmasked model-to-map FSC was calculated in PHENIX for the final model against the reconstructed map.

## Acknowledgements

S.L. thanks Tsinghua University for providing a Start-up fund, the Tsinghua University Branch of China National Center for Protein Sciences (Beijing) for the cryo-EM facility and the computational facility support, and Dr. Fan Yang, Liao Zhang for technical support. We thank Dr. Jianlin Lei in the cryo-EM Facility, Dr. Lingpeng Cheng in the Cryo-EM Structural Analysis Facility, and Dr. Haiteng Deng in the Proteinomics Facility at Technology Center for Protein Sciences, Tsinghua University, for cryo-EM data collection, structural analysis and protein MS; Haibo Wu, Changzhong Jin and Zhigang Wu in the State Key Laboratory for Diagnosis and Treatment of Infectious Diseases, The First Affiliated Hospital, Zhejiang University School of Medicine, for virus fixation and biosafety validation. We thank the computational facility support on the cluster of Bio-Computing Platform (Tsinghua University Branch of China National Center for Protein Sciences Beijing). We are in debt to Dr. Hongwei Wang and Dr. Nieng Yan for providing critical advices.

## Author contributions

S.L. conceptualized the models and supervised the project. Shibo L. isolated virus and conducted clinical diagnosis. H.Y., N.W., M.Z., D.S. and X.L. propagated and fixed the virus sample. YS purified the virus, prepared EM grids and collected cryo-ET data. Y.S., Z.Z., C.P., W.K., J.W., Y.C. and S.L. analyzed cryo-ET data. J.X., Y.S. and M.E.G. collected the cryo-EM data. J.X. analyzed the cryo-EM data and built the model. Z.Z. and Y.S. performed the statistical analysis. Y.S. and C.Z. collected and analyzed MS data. S.L. wrote the original draft. W.K., C.P., Y.S., H.Y. and S.L. reviewed and edited the manuscript. Y.S., Z.Z., J.X. and C.P. prepared the figures. S.L., H.Y. and N.W. acquired funding and administrated the protect.

## Declare of interests

The authors declare no competing interests.

