## Supplementary information for "In situ architecture and membrane fusion of SARS-CoV-2 Delta variant"

**The Supplementary material includes:**

Supplementary Movie 1

Supplementary Figures 1 to 8

Supplementary Tables 1 to 2

**Supplementary Movie 1.** Cryo-electron tomogram as shown in Figure 1a.

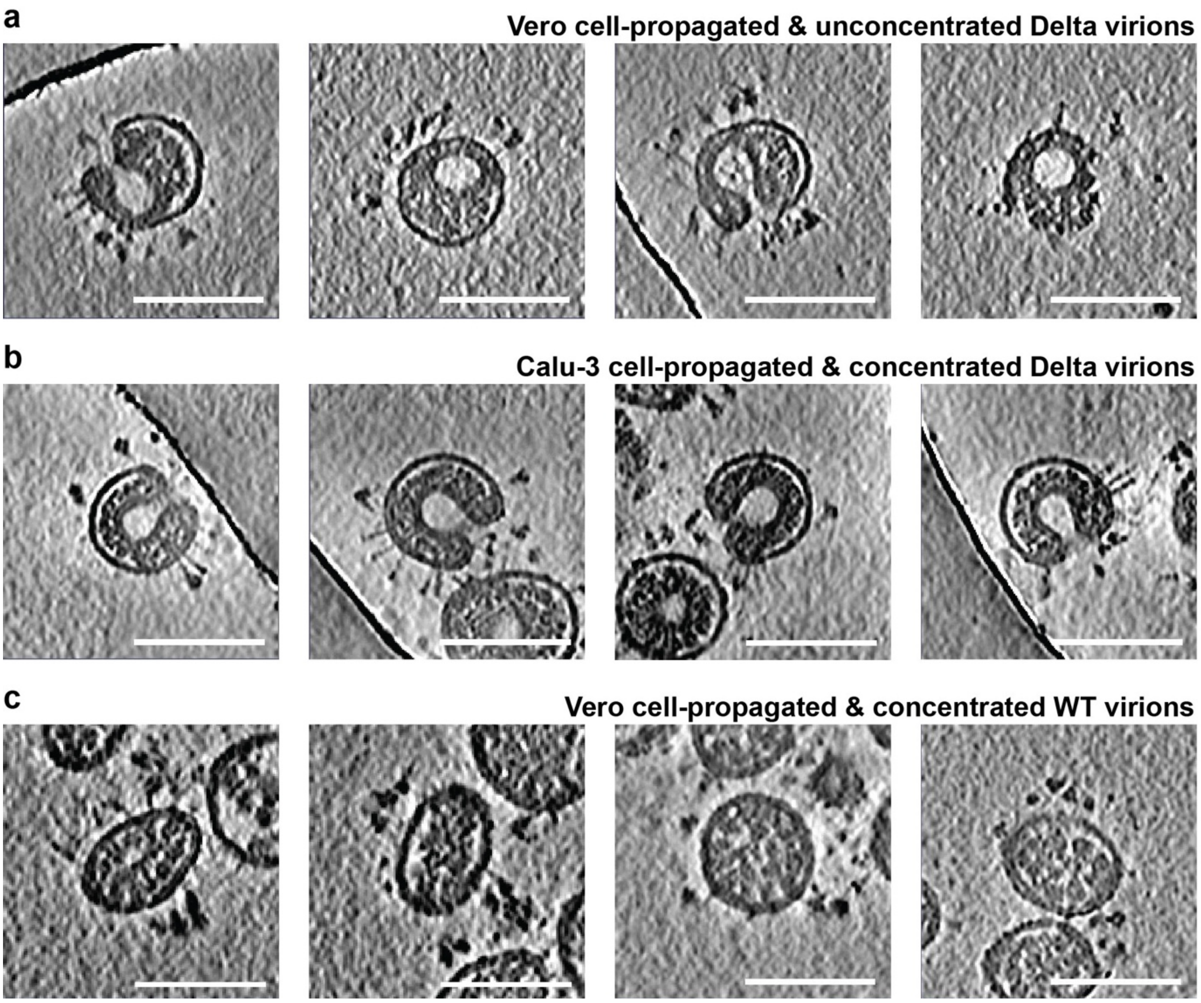


**Supplementary Figure 1. Validations of SARS-CoV-2 Delta virion invagination in various conditions.**

Cryo-electron tomography slices (5 nm thick) of (**a**) Delta virions in the supernatant of infected Vero cells. (**b**) Calu-3 cell-propagated Delta virions concentrated onto the interface between 30% and 50% sucrose cushion by ultracentrifugation. Both conditions contain invaginated virions. (**c**) Vero cell-propagated WT virions concentrated onto the interface between 30% and 50% sucrose cushion by ultracentrifugation. Invaginated virions were rarely seen in this condition. Scale bars: 100 nm.

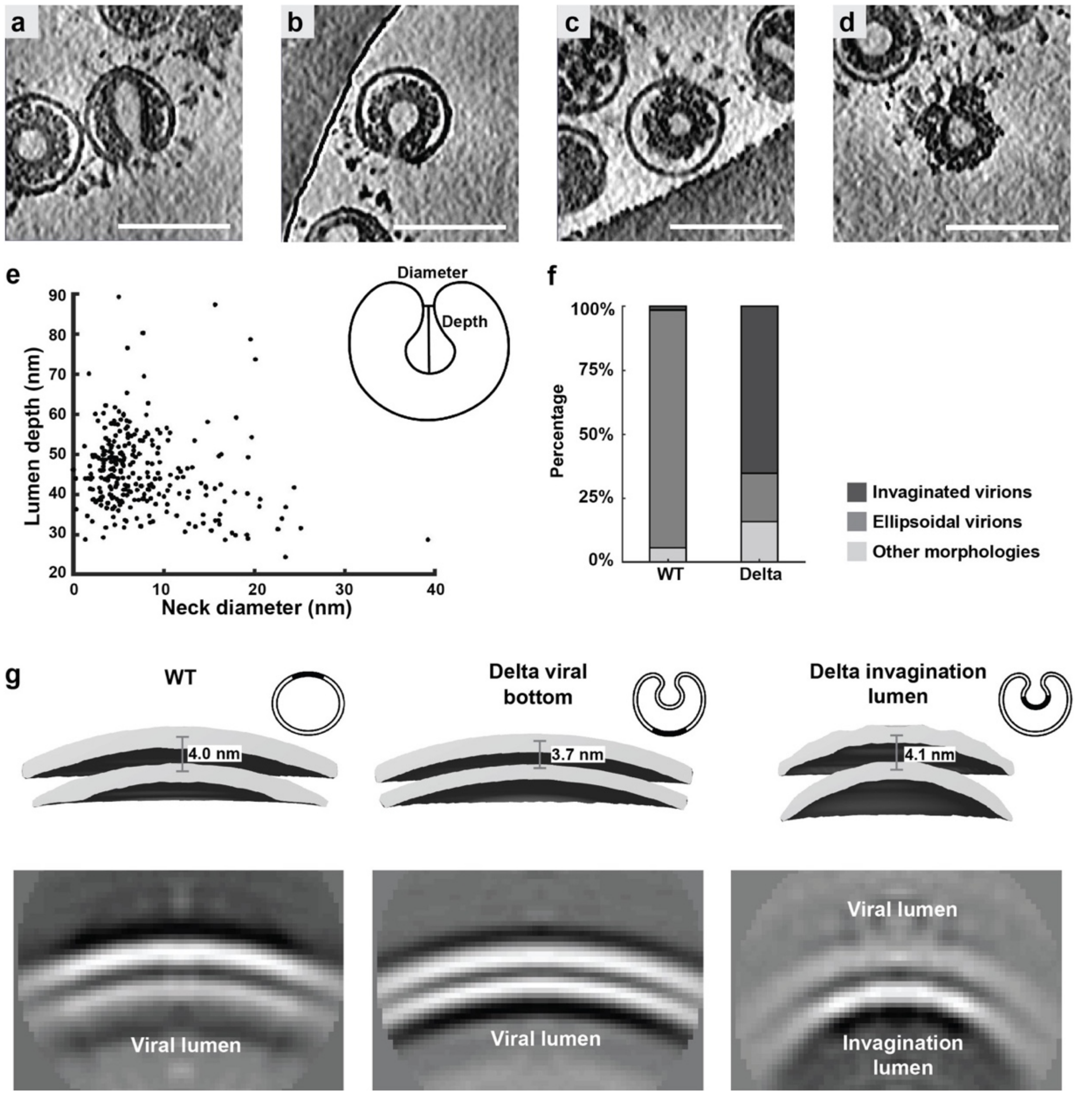


**Supplementary Figure 2. Characterization of the invaginated SARS-CoV-2 Delta virions.** (**a-d**) Tomogram slices (5 nm thick) showing typical features of invaginated virions, including (a) clustering of S around the neck area; (b) the majority of RNPs wrapping around the invagination, leaving the rest of the viral lumen empty; (c) S existing only in areas where RNPs are attached to the cytoplasmic side of the envelope and that (d) the viral bottom often appears broken. Scale bars: 100 nm. (**e**) Statistics of the invagination depth and neck diameter. (**f**) Percentage of invaginated, ellipsoidal (spherical) and other virion morphologies observed in the WT and Delta virion population. (**g**) Comparison of the thickness among envelopes from SARS-CoV-2 WT virions, Delta viral bottom and Delta invagination lumen. Subtomogram averages of the lipid-bilayer densities were shown as 3D reconstruction at the same threshold (top) and as 2D slices (bottom).

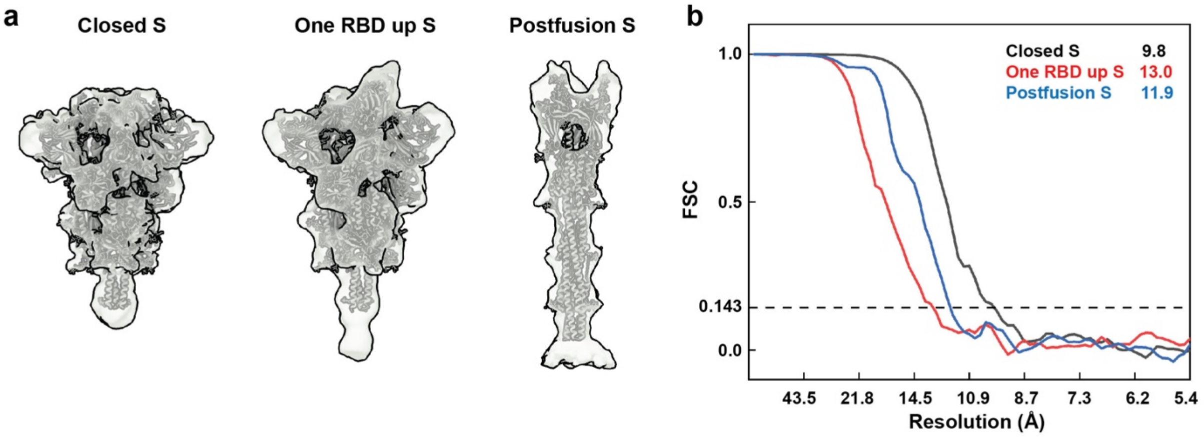


**Supplementary Figure 3. Local resolution and Fourier Shell Correlation (FSC) curves of structures solved by subtomogram averaging.**

(**a**) Maps of the closed and one RBD up prefusion S and postfusion S were fitted with corresponding recombinant structures (PDB:7SBK, 7SBL and 6XRA, relatively) for comparison. (**b**) Resolutions of closed, one RBD up and postfusion S were estimated form FSC curves, using a criterion of 0.143.

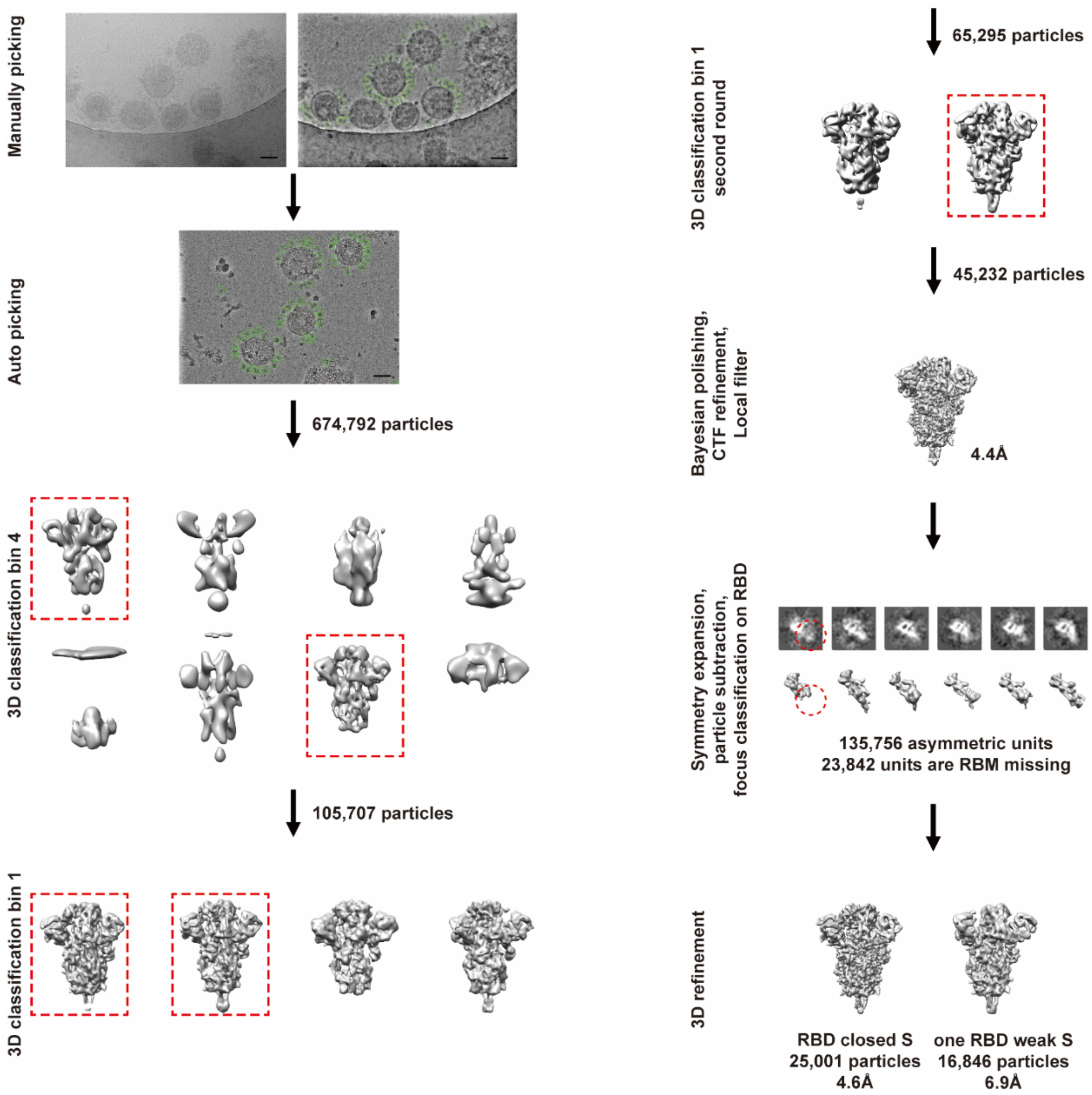


**Supplementary Figure 4. Single particle cryo-EM image processing workflow.**
Initial micrographs were deconvolved to allow automatic particle picking (green circles) and 3D classification. Scale bar, 50 nm. Selected classes indicated with red dashed boxes were refined to a combined RBD closed S at 4.4 Å resolution. Symmetry expansion was applied on the class and RBDs were subtracted for local classification to sort different conformations. The locally classified asymmetric units are shown in topview, which differ in density weakness. The RBD closed S were refined with C3 symmetry applied. The one RBD weak S were refined with C1 symmetry applied. For further details see materials and methods.

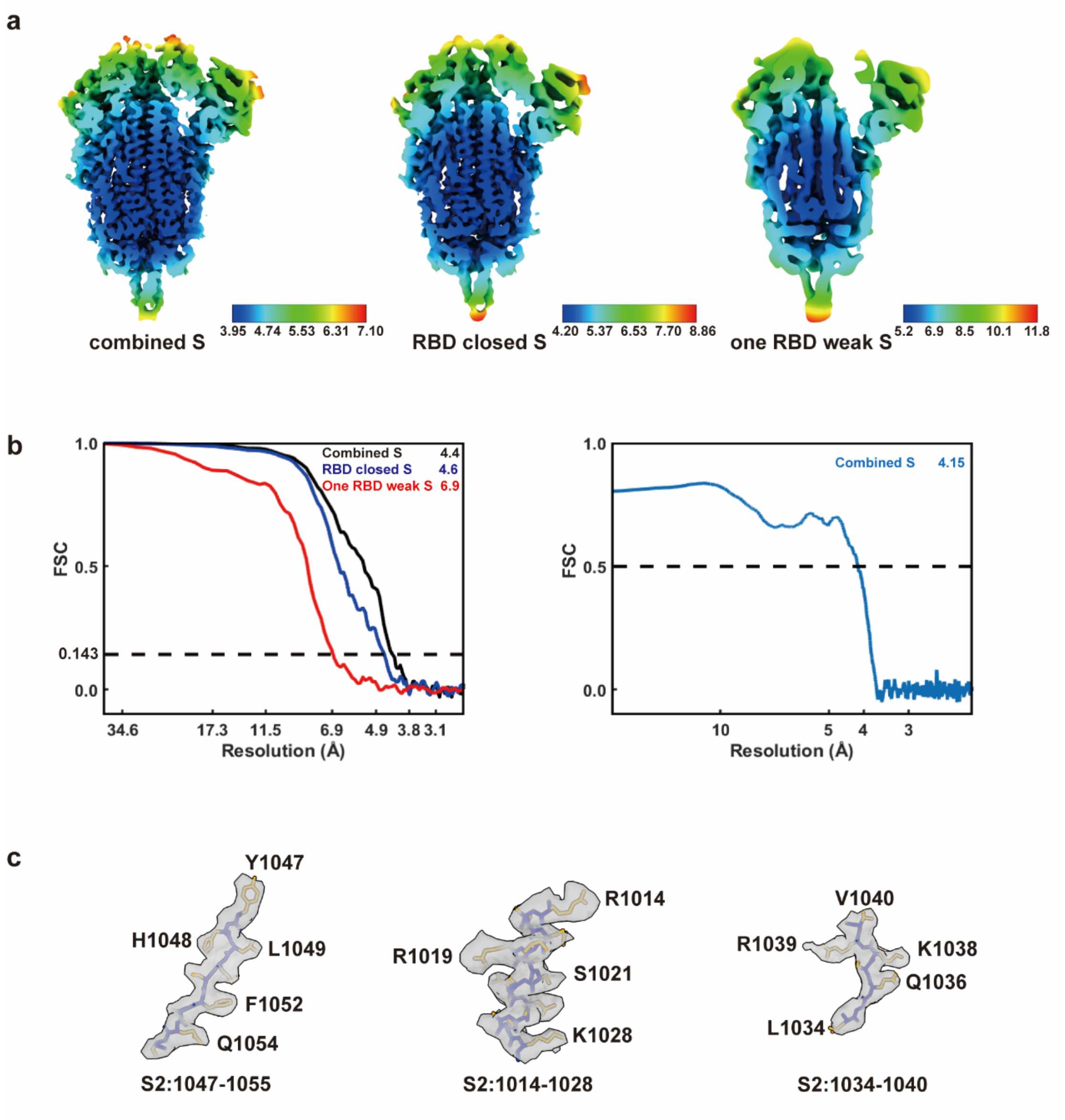


**Supplementary Figure 5. Local resolution and Fourier Shell Correlation (FSC) curve of the native closed prefusion S structure solved by single particle analysis.**

(**a**) Maps of the combined, closed and one RBD weak S colored by their local resolution. (**b**) FSC curves of three structures (left, resolution was estimated using a criterion 0.143) and for the atomic model against the map (right, resolution was estimated using a criterion 0.5). (**c**) The best solved peptides of the combined closed S, which locate on S2: 1047-1055, 1014-1028, and 1034-1040, were highlighted with the refined model.

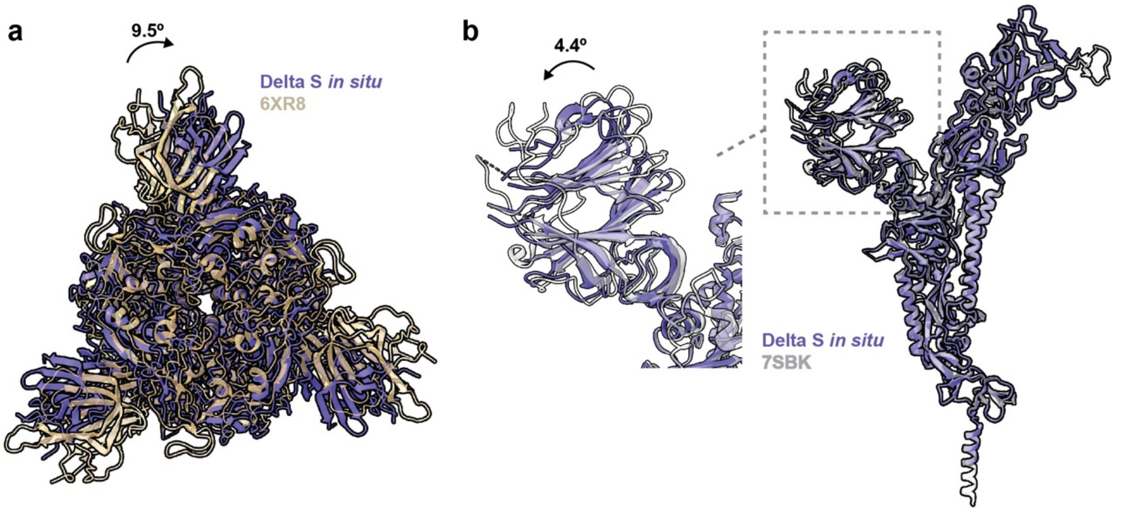


**Supplementary Figure 6. Comparison between the structures of native SARS-CoV-2 Delta S and recombinant WT S in the closed conformation.**

(**a**) The in situ structure of Delta S was fitted with that of the recombinant WT S (PDB:6XR8), showing a 9.9° clockwise rotation of NTD on the native Delta S. (**b**) Superposition of the in situ Delta S and the recombinant Delta S (PDB:7SBK), showing the NTD in our structure extends 4.4° outward when S2 of both structures are aligned.

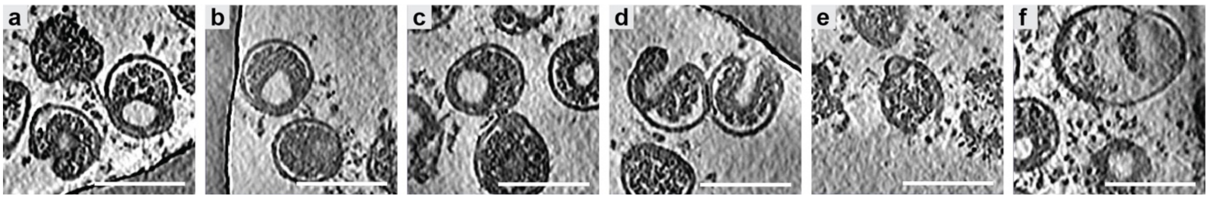


**Supplementary Figure 7. Further representative tomogram slices and quantitative descriptions of six membrane fusion steps of SARS-CoV-2 Delta virions.**

A tomogram slice of (**a**) bridging, (**b**) dimpling, (**c**) pinching, (**d**) tightly docked interface, (**e**) hemifusion and (**f**) fusion completion. Tomogram slice thickness: 5 nm; Scale bars: 100 nm.

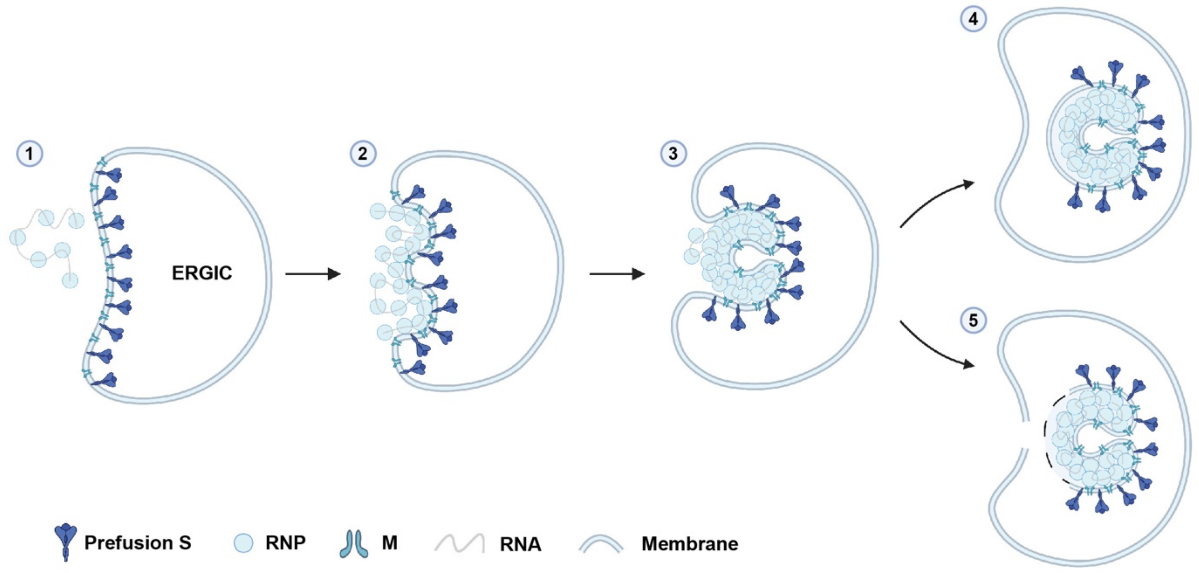


**Supplementary Figure 8. Proposed model for the invagination formation in Delta variant.** In this model, N-mutations have changed the assembly and RNA-packaging of N, resulting in an altered mode of viral budding into the ER-Golgi intermediate compartment (ERGIC) to produce progeny virions. (1) RNPs are recruited by M residing on the membrane of an EGRIC, which harbors S-trimers on the opposite of the membrane. (2) RNPs bind with M and bud into the ERGIC while forming a dimple. (3) When budding proceed, the dimple enlarges into an invagination. With increased budding efficiency or change of RNP-assembly due to N-mutations, the viral bottom was left either (4) bald or (5) broken after budding completion. Model was created with BioRender.

**Supplementary Table 1. Cryo-ET data acquisition and reconstruction statistics.**

| **Data collection** | | | |
|---|---|---|---|
| Microscope | Titan Krios | | |
| Magnification | 64,000 | | |
| Voltage (kV) | 300 | | |
| Detector | Gatan K3 | | |
| Energy filter | Gatan GIF Quantum, 20 eV slit | | |
| Pixel size (Å) | 0.68 (super-resolution) | | |
| Tilt schemes | Dose-symmetric scheme | | |
| Number of tilt-series | 150 | | |
| Number of virions | 1,032 | | |
| Exposure ($e^-/Å^2$) | 131.2 | | |
| Defocus range (µm) | -2.0 ~ -4.0 | | |
| Software | SerialEM | | |
| **Reconstruction** | | | |
| Software | Dynamo 1.1.333 | | |
| Dataset | Prefusion S (Closed) | Prefusion S (One RBD up) | Postfusion S |
| Final number of particles | 8,269 | 14,426 | 2,810 |
| Symmetry imposed | C3 | C1 | C3 |
| Final Resolution (Å) | 9.8 | 13.0 | 11.9 |
| Gold-standard | yes | yes | yes |
| FSC threshold | 0.143 | 0.143 | 0.143 |
| Final pixelsize (Å) | 2.72 | 2.72 | 2.72 |
| Map sharpening B-factor ($Å^2$) | -1500 ~ -2000 | N/A | N/A |

**Supplementary Table 2. Cryo-EM data acquisition, refinement and validation statistics.**

| | | |
|---|---|---|
| **Data collection** | | |
| Microscope | Titan Krios | |
| Magnification | 81,000 | |
| Voltage (kV) | 300 | |
| Detector | Gatan K3 | |
| Energy filter | Gatan GIF Quantum, 20 eV slit | |
| Pixel size (Å) | 0.541 (super-resolution) | |
| Movies collected | 25,851 | |
| Movies for final reconstruction | 18,028 | |
| Exposure ($e^-/Å^2$) | 50 | |
| Defocus range (μm) | -1.0 ~ -3.0 | |
| Software | AutoEmation 2 | |
| **Reconstruction** | | |
| Software | Relion 3.1.0 | |
| Symmetry imposed | C3 | |
| Initial particle numbers | 674,792 | |
| Final particle numbers | 45,232 | |
| Final map resolution (Å) | 4.39 | |
| FSC threshold | 0.143 | |
| Final pixelsize (Å) | 1.082 | |
| **Classification** | | |
| Conformations | Three RBDs down | Three RBDs down with one RBD weak |
| Particle numbers | 25,001 | 16,846 |
| Symmetry imposed | C3 | C1 |
| Map resolution (Å) | 4.61 | 6.92 |
| **Refinement** | | |
| Initial model used | PDB 6XR8 & 7SBK | |
| Model resolution (Å) | 4.1 | |
| FSC threshold | 0.5 | |
| Map sharpening *B* factors ($Å^2$) | -200 | |
| Model composition | | |
| Non-hydrogen atoms | 24,684 | |
| Protein residues | 3,036 | |
| Ligands | 66 | |
| *B* factors ($Å^2$) | | |
| Protein | 118.79 | |
| Ligand | 106.67 | |
| R.m.s deviations | | |
| Bond lengths (Å) | 0.007 | |
| Bond angels (°) | 1.417 | |
| **Validation** | | |
| MolProbity score | 1.74 | |
| Clashscore | 4.78 | |
| Poor rotamers (%) | 1.88 | |
| Ramachandran plot | | |
| Favored (%) | 95.88 | |
| Allowed (%) | 4.02 | |
| Disallowed (%) | 0.10 | |
